# Soil bacterium *Massilia* secretes metabolites that promote *Leptospira* growth

**DOI:** 10.64898/2026.04.06.716759

**Authors:** Michinobu Yoshimura, Ryo Ozuru, Satoshi Miyahara, Fumiko Obata, Mitsumasa Saito, Takumi Sonoda, Yusuke Kurihara, Jason A. Papin, Glynis L. Kolling, Shin-ichi Yoshida, Kenji Hiromatsu

**Author notes:** Address co-correspondence to Michinobu Yoshimura. Address co-correspondence to Ryo Ozuru. Department of Pathology, Brigham and Women’s Hospital, Boston, Massachusetts, USA. Department of Parasitology, Kurume University, Kurume, Fukuoka, Japan (Yusuke Kurihara).

## Abstract

Understanding pathogen metabolism is critical for identifying key functions for drug targeting, establishing effective *in vitro* experimental systems, etc., particularly for metabolically unique organisms such as *Leptospira*. Pathogenic *Leptospira* are thought to infect humans from environmental sources; however, direct isolation from environmental samples remains technically challenging and is not yet well established. Here, we report that a ubiquitous environmental bacterium, *Massilia* sp., produces metabolites that promote the growth of *Leptospira interrogans*, encountered through an incidental contamination event, and identified in this study. Gas chromatography–tandem mass spectrometry (GC-MS/MS) analysis showed demonstrated that cultivating of *Massilia* sp. in R2A medium resulted in the accumulation of metabolites, including branched-chain amino acid (BCAA) intermediates, compared to fresh medium. By combining genome-scale metabolic modeling with experimental validation using cell-free culture supernatant supplementation assays, we demonstrate that BCAA intermediates, particularly 2-ketoisocaproic acid (4-methyl-2-oxopentanoate; 4MOP), a leucine biosynthetic intermediate produced by *Massilia* sp., enhance *Leptospira* growth. To investigate the metabolic role of 4MOP, we incorporated transcriptomic data into a genome-scale metabolic network model to generate condition-specific models. Resulted flux distributions indicated that *Leptospira* catabolized imported 4MOP to produce acetyl-CoA. Our results reveal a previously unrecognized metabolic interaction where metabolites produced by environmental bacteria support the growth of pathogenic *Leptospira*, offering mechanistic insight into its metabolic requirement. These findings have implications to understand the environmental persistence of *Leptospira* through its metabolic dependencies on coexisting microbes, and they also help develop better strategies for this pathogen.

**Importance:** Pathogenic *Leptospira* persist in environmental reservoirs, yet the mechanisms supporting their growth remain poorly defined. Here, we find that metabolites produced by common environmental bacteria, *Massilia* sp., can promote *Leptospira* growth, suggesting a previously unrecognized metabolic dependency on coexisting microbes. Importantly, this study indicates that combining genome-scale metabolic modeling with experimental validation provides a useful framework for identifying metabolic interactions that are otherwise difficult to resolve using conventional culture-based approaches. Current strategy may facilitate the systematic identification of growth-supporting metabolites and provide a basis for improving selective cultivation for uncultured or difficult to culture organisms. Determination of growth promoting metabolites advances our understanding of pathogen persistence in natural environments and offers a generalized framework to study metabolically dependent microorganisms.

## Introduction

Zoonotic pathogens present a serious One Health challenge (1). After being shed from infected hosts, many bacteria persist in environmental reservoirs such as soil and surface water, where they can trigger new outbreaks (2). Yet the mechanisms that support survival and, in some cases, proliferation outside the hosts remain poorly understood, especially under limited nutrient condition, physicochemical stress, and microbial competition. Because soil and surface water harbor dense and metabolically active microbial communities, pathogens are inevitably embedded in a chemically dynamic milieu shaped by microbial exudates (3). Environmental bacteria can enrich microenvironments with growth-supporting molecules including diffusible metabolites that pathogens may be unable to synthesize at all or able to internalize inefficiently from exogenous sources (4, 5). We therefore hypothesized that secreted products from environmental bacteria can promote the growth of zoonotic pathogens, and that identifying these products will improve mechanistic understanding of pathogen maintenance in environmental reservoirs.

*Leptospira* spp., the spirochete responsible for leptospirosis, offer an experimentally accessible system to study how environmental microbial products can influence the ecology of zoonotic pathogens. Leptospirosis is one of the most widespread zoonoses worldwide (6, 7). Rodents, especially the genus *Rattus*, serve as principal reservoirs, harbor *Leptospira* in renal tubules, and shed bacteria in urine. Once released, the organisms spread through surface water and soil, where they can infect wildlife, livestock, companion animals, and humans via cutaneous or mucosal exposure (8). In humans, infection often appears as a self-limited influenza-like illness that responds to prompt antibiotic treatment, yet a subset of patients progresses to severe disease, including Weil’s disease characterized by jaundice, pulmonary hemorrhage, and acute kidney injury, with a significantly increased risk of death (9). In animals, clinical outcomes vary by host species, and severe cases can cause abortions and stillbirths, imposing major economic burdens on livestock industries (10, 11). These features exemplify the interconnected relationships among human, animal, and environmental dimensions that are central to One Health. Notably, *Leptospira* grows very slow *in vitro* and requires nutritionally complex media (12) suggesting that its growth may be limited by restricted access to key metabolites in environmental reservoirs. Such potential dependency led to the hypothesis that diffusible products released by co-occurring environmental bacteria can provide growth supporting factors for *Leptospira*. Identifying such factors will clarify mechanisms that contribute to pathogen maintenance in soil and surface waters.

During routine cultivation, we serendipitously observed that a co-isolated soil bacterium, *Massilia* sp., significantly accelerates *Leptospira* proliferation (Fig S1). Motivated by this finding, we aimed to identify the *Massilia*-derived diffusible factors that promote *Leptospira* growth, and to define the *Leptospira* metabolic processes that enable their utilization. To this end, we implemented an integrated systems biology workflow combining metabolomic profiling of *Massilia* culture supernatants with genome-scale metabolic network reconstruction (GENRE) (13, 14). Metabolomics, together with *in silico* biomass simulations, prioritized candidate growth-promoting metabolites, which we subsequently validated *in vitro*. We then used contextualized (*i.e.* transcriptome-informed) GENREs to delineate the metabolic routes through which *Leptospira* assimilates these compounds under proliferative conditions. Collectively, this framework links an environmental bacterial exometabolome with mechanistic predictions and experimental validations in nutrient-limited pathogen growth.

## Results

### *Massilia*-conditioned supernatant enhances *Leptospira* growth yield

To determine whether diffusible products released by *Massilia* can promote *Leptospira* growth, we measured the effect of cell-free *Massilia* culture supernatant (Msup) on *Leptospira* growth curves. Adding Msup to the culture medium with Msup consistently increased the overall growth yield of *L. interrogans* compared to control conditions, as shown by a higher terminal optical density (Fig. 1A and B). Notably, the primary effect was on growth yield rather than the apparent growth rate (Fig. S2). This effect was not restricted to a single *Leptospira* strain. Supernatant from a representative *Massilia* sp. enhanced the growth of avirulent (P2 clade) and saprophytic (S clade) *Leptospira* species (Fig. 1C and D), indicating a broad effect across the genus with a few exceptions (E8, P2 clade). Together, these results indicate that *Massilia* releases extracellular factors that promote growth and increase *Leptospira* biomass, leading to further identification of the active components within the secreted mixture (exometabolites).

**Fig 1.**
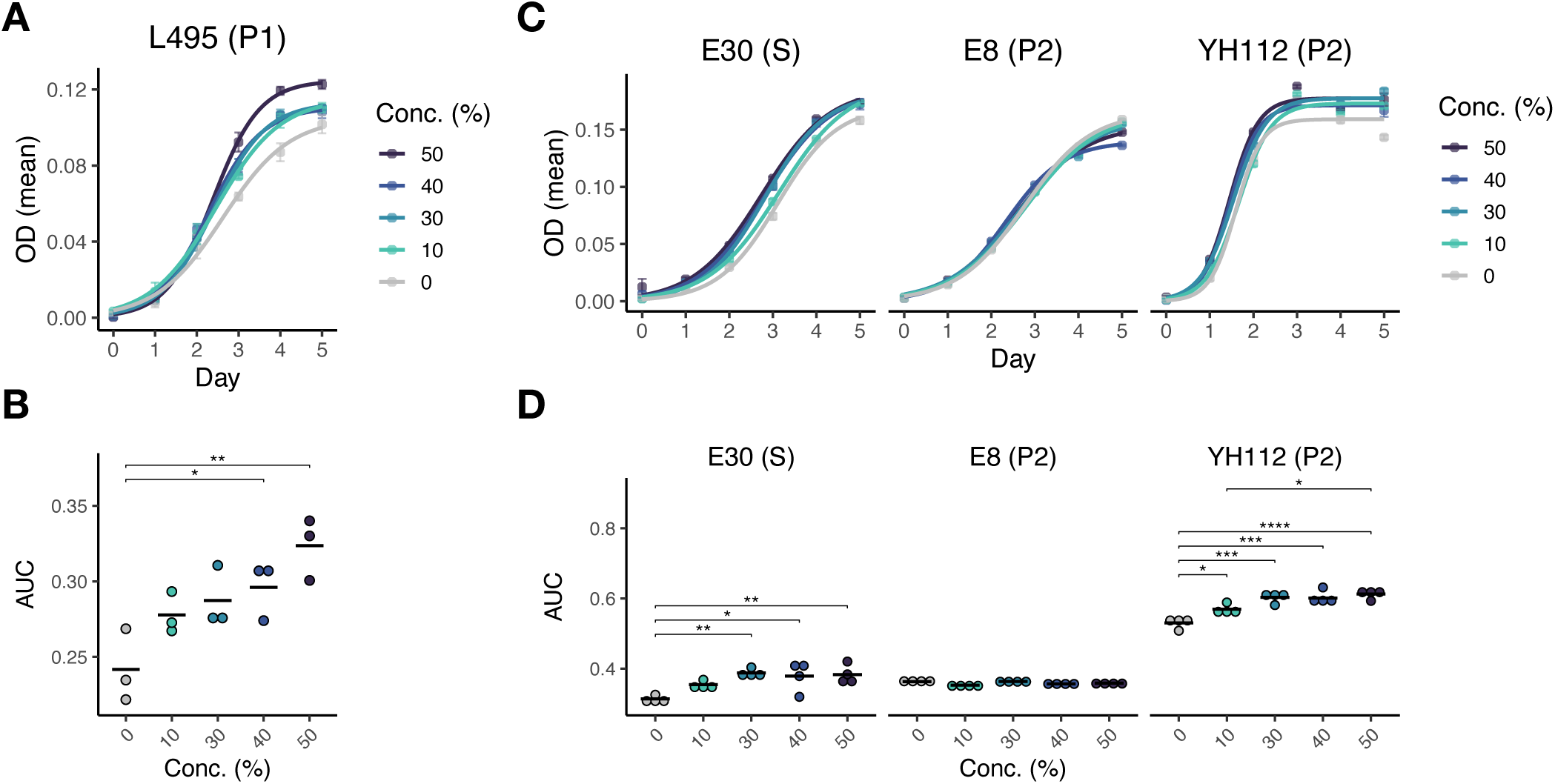
Growth-promoting effect of Msup on *Leptospira*. (A) Growth curve of *Leptospira interrogans* serovar Manilae strain L495 (pathogenic clade: P1) following supplementation with *Massilia* culture supernatant (Msup). Curves represent the mean of triplicate cultures (n = 3). (B) Area under the curve (AUC) calculated from the growth curves shown in (A). (C) Growth curves of multiple *Leptospira* strains (avirulent strain: P2 and saprophytic strain: S) supplemented with Msup. Curves represent the mean of quadruplicate cultures (n = 4). (D) Area under the curve (AUC) calculated from the growth curves shown in (C). Growth was monitored by measuring OD_450_. Statistical significance was evaluated using one-way ANOVA followed by Tukey’s HSD post hoc test. *P < 0.05, **p < 0.01, ***p < 0.001, and ****p < 0.0001

### Metabolomics and GENRE prioritize candidate growth-promoting metabolites derived from Massilia

To identify diffusible metabolites in Msup that account for the biomass-increasing activity, we first profiled Msup using gas chromatography–tandem mass spectrometry (GC-MS/MS)-based metabolomics, and then evaluated candidate compounds with genome-scale metabolic network reconstruction (GENRE) (Fig. 2). We first analyzed the chemical composition of cell-free *Massilia* culture supernatant and the corresponding fresh R2A medium control using GC-MS/MS. Principal coordinate analysis (PCoA) revealed that Msup had a composition different from R2A alone (Fig. 2A). We then defined a set of metabolites characteristic of Msup that is significantly enriched in Msup relative to R2A (FDR adjusted t test p value < 0.05) or detected exclusively in Msup. These candidates are summarized in Fig. 2B, with individual samples labelled R1 to R3 for R2A controls and M1 to M4 for Msup replicates. Among the prioritized metabolites, glycine was the only proteinogenic amino acid identified as a single-amino-acid species.

**Fig 2.**
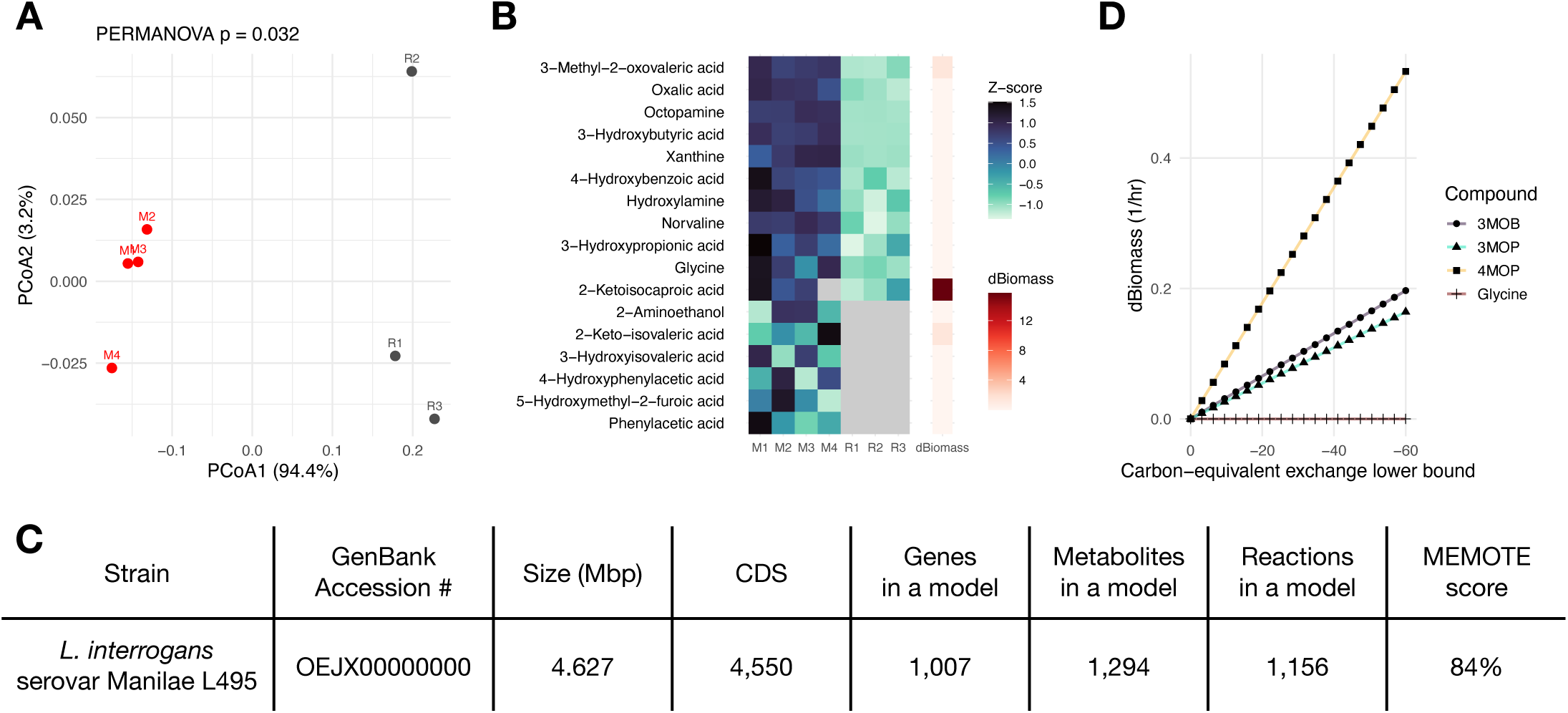
Metabolomic profiling of Msup and model-guided prioritization of candidate growth-promoting compounds. (A) Principal coordinates analysis (PCoA) of GC-MS/MS-based metabolomic profiles comparing Msup (M1-M4) with medium-only controls (R1-R3). (B) Heatmap of metabolites characteristically detected in Msup, in which each compound is color-coded according to z-score. The predicted effects of each compound on Leptospira proliferation based on metabolic simulations, were expressed as changes in biomass production (dBiomass). Gray indicates signals below the detection limit. (C) Genomic data and the model characteristics in this study. (D) In silico growth simulation results showing dBiomass relative to the base medium control after supplementation with candidate compounds across a range of carbon-equivalent exchange lower bounds (C-mmol gDW⁻¹ h⁻¹). Because negative exchange flux denotes substrate uptake in this model, more negative x-axis values indicate greater substrate availability. 4MOP; 4-methyl-2-oxopentanoate, 3MOP; 3-methyl-2-oxopentanoate, 3MOB; 3-methyl-2-oxobutanoate.

Because metabolomics alone generates a diverse set of candidates, we then used GENRE to prioritize metabolites predicted to directly support *Leptospira* biomass formation. Draft GENREs for pathogenic *Leptospira interrogans* strain L495 used in this study were generated with Reconstructor (15). Culture conditions were represented as Ellighausen-McCullough-Johnson-Harris liquid medium (EMJH, Table S1). Genome references and model statistics including gene counts, metabolites, reactions, and MEMOTE scores (16) are provided in Fig. 2C. Using the *Leptospira* model, we performed *in silico* supplementation by allowing uptake of each Msup-associated metabolite and quantifying the resulting change in predicted biomass production relative to the base medium control (Fig. 2B; dBiomass). Among the tested metabolites, 2-Ketoisocaproic acid, also known as 4-methyl-2-oxopentanoate (4MOP), a branched-chain amino acid (BCAA) intermediate in leucine catabolism, produced the largest increase in biomass (Fig. 2B; dBiomass). Because the GC-MS/MS dataset also contained additional BCAA intermediates that increased dBiomass, including 3-Methyl-2-oxovaleric acid and 2-keto-isovaleric acid, also known as 3-methyl-2-oxopentanoate (3MOP) and 3-methyl-2-oxobutanoate (3MOB), respectively, we extended the simulations to include these compounds as well as glycine. Varying the carbon-equivalent exchange lower bound for each candidate compound, it showed a robust increase in biomass production with 4MOP, whereas the other tested metabolites had limited or no effect under the modeled conditions (Fig. 2D). This indicates an agreement between detected metabolites and simulated dBiomass.

### BCAA intermediates, including 4MOP, enhance *Leptospira* growth

To experimentally evaluate the GENRE-based based prioritization of BCAA intermediates as candidate stimulatory metabolites, we supplemented EMJH with either the BCAAs or their corresponding keto acid intermediates (4MOP, 3MOP, and 3MOB) (Fig. 3A and B). Consistent with the model-based prediction that BCAA intermediates can support biomass formation, 4MOP promoted *Leptospira* growth; however, 3MOP, 3MOB, and mixtures of these intermediates also increased growth yield, indicating that growth promotion is not specific to 4MOP alone but extends to multiple BCAA-derived intermediates under the tested conditions. According to the result, the highest concentration (10 µM) tested for 4MOP and 3MOP restricted growth below the non-supplemental condition, which suggests there may be an optimized condition and a fine-tuning mechanism within *Leptospira*.

**Fig 3.**
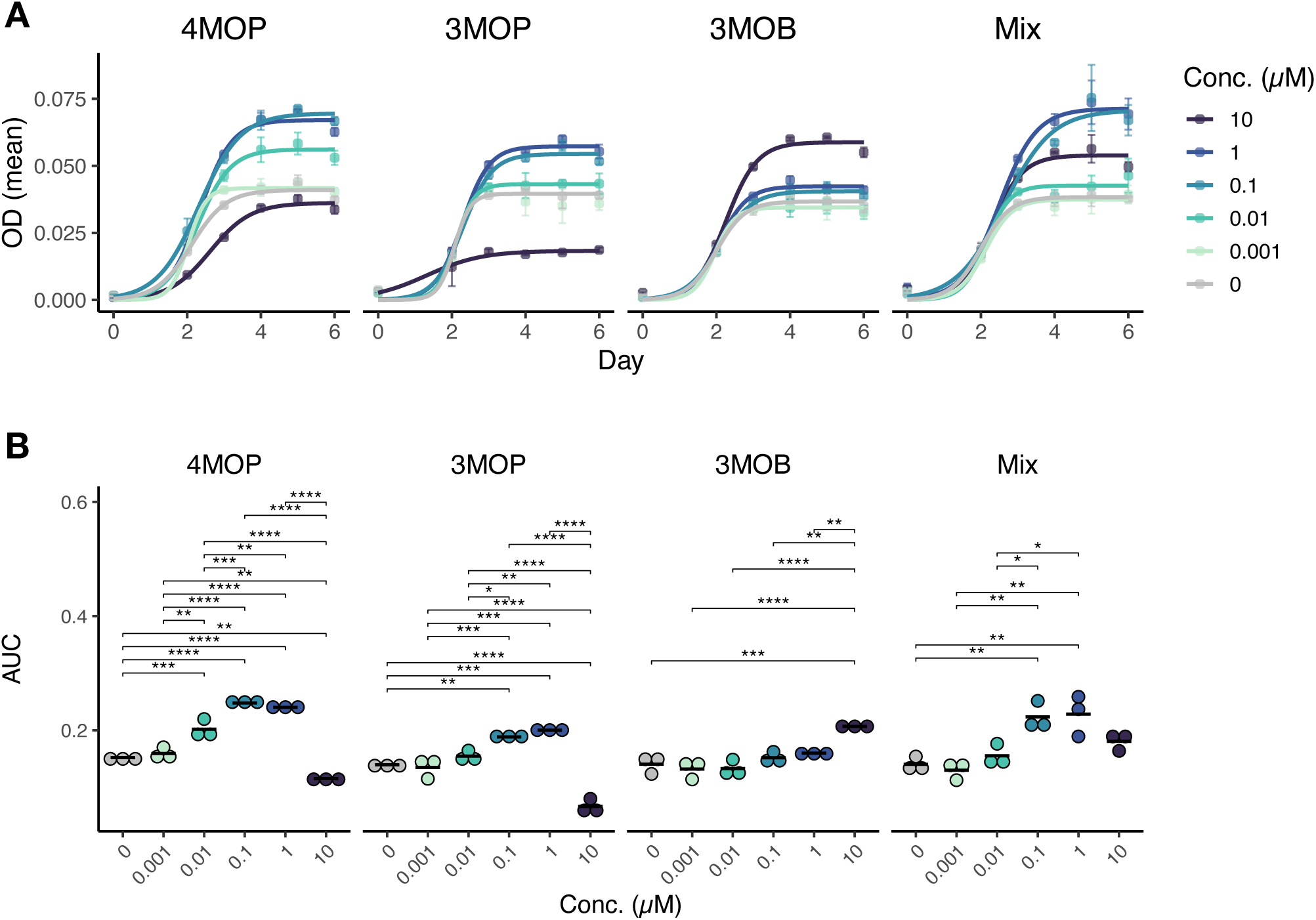
Effects of branched-chain amino acid (BCAA) intermediates on *Leptospira* growth. (A) Growth curves of *Leptospira* following supplementation with BCAA intermediates: 4-methyl-2-oxopentanoate (4MOP), 3-methyl-2-oxopentanoate (3MOP), 3-methyl-2-oxobutanoate (3MOB), or a mixture of these intermediates (Mix). Curves represent the mean of triplicate cultures (n = 3). (B) Area under the curve (AUC) calculated from the growth curves shown in (A). Growth was monitored by measuring OD_450_. Statistical significance was evaluated using one-way ANOVA followed by Tukey’s HSD post hoc test. *P < 0.05, **p < 0.01, ***p < 0.001, and ****p < 0.0001

### Contextualized GENRE implicates an increase in flux through leucine catabolism under the influence of Msup

To gain mechanistic insight into how Msup supports *Leptospira* growth, we generated a contextualized GENRE for *L. interrogans* strain L495 (Fig. 4A). L495 was cultured under three conditions, EMJH alone, EMJH supplemented with R2A, and EMJH supplemented with Msup. Bacterial RNA was collected 48 hours post inoculation for transcriptome profiling. Transcript abundances were integrated into the L495 metabolic model using RIPTiDe (17), which constrains and weights feasible flux distributions based on transcriptome data. We interrogated BCAA-associated pathways highlighted by the metabolomics-guided GENRE prioritizations and supplementation simulation. Flux patterns in the Msup-contextualized model indicated an increase in utilization of the leucine degradation pathway, including reactions that convert BCAA-derived keto-acid intermediates toward acetyl-CoA generating routes (Fig. 4B). For example, rxn38054 catalyzes the conversion of leucine to 4MOP, and its increased flux under the Msup condition predicts enhanced metabolic conversion from leucine to 4MOP. Downstream of this step, the predicted flux was connected to the production of acetoacetate and acetyl-CoA. Acetoacetate is further converted to acetoacetyl-CoA and subsequently to acetyl-CoA, suggesting that incoming 4MOP can be efficiently funneled into acetyl-CoA production in the model. These changes provide a systems-level explanation for the observed growth-supporting activity of BCAA intermediates, linking Msup-associated transcriptional states to enhanced capacity for carbon assimilation through leucine catabolism.

**Fig 4.**
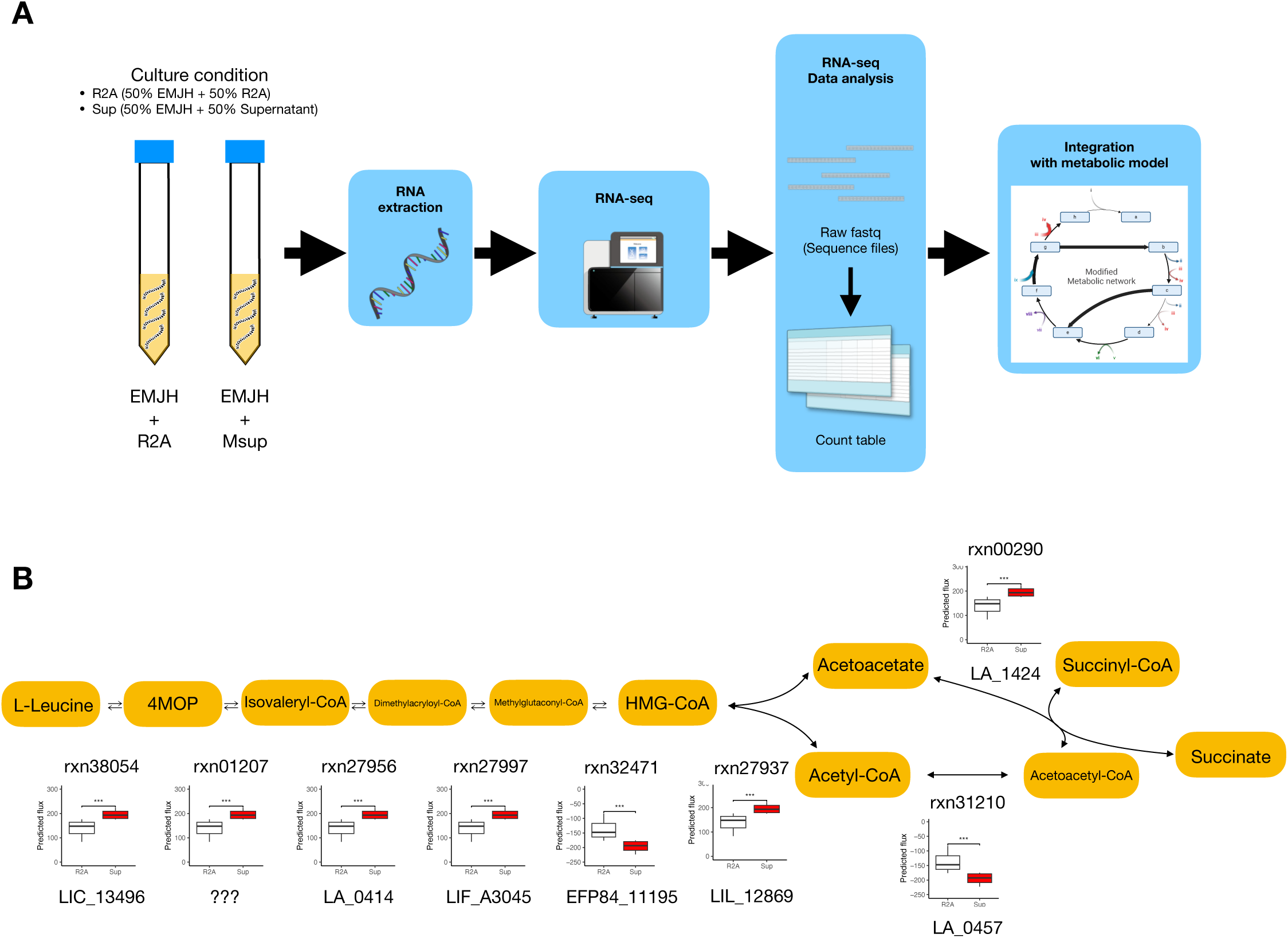
Context-specific GENRE analysis reveals *Leptospira* metabolic changes upon supplementation with Msup. (A) Experimental overview and definition of *Leptospira* culture conditions. (B) Predicted changes in sampled fluxes within the leucine degradation pathway upon Msup supplementation compared with the R2A condition. For each box plot panel, the ModelSEED reaction ID is shown above, and candidate *Leptospira* genes annotated by Reconstructor for the corresponding reaction are shown below. Reactions proceeding to the right are defined as the forward direction, except for rxn32471, that the corresponding gene EFP84_11195 contributes to convert Acetyl-CoA to HMG-CoA. The gene for rxn01207 was not represented in the metabolic model analyzed in this study. Statistical significance was evaluated using the Wilcoxon rank-sum test. *P < 0.05, **p < 0.01, ***p < 0.001, and ****p < 0.0001

## Discussion

This study demonstrates that cell-free culture supernatant from the soil bacterium *Massilia* increases *Leptospira* growth yield. By integrating GC-MS/MS-based metabolomics with GENRE, we prioritized candidate metabolites present in the *Massilia*-conditioned media and identified BCAA-derived keto acid intermediates, such as 4MOP, as plausible growth-promoting compounds (Fig. 5). Supplementation experiments confirmed this prioritization by showing that these intermediates boost *Leptospira* growth yield *in vitro*. To place these observations in a mechanistic context, transcriptome-informed model contextualization using RIPTiDe (17) was consistent with a metabolic shift toward increased utilization of leucine catabolic routes. Collectively, our results illustrate how an initial phenotypic observation can be advanced to candidate identification and testable mechanistic hypotheses through the integration of multi-omics data and systems-level modelling.

**Fig 5.**
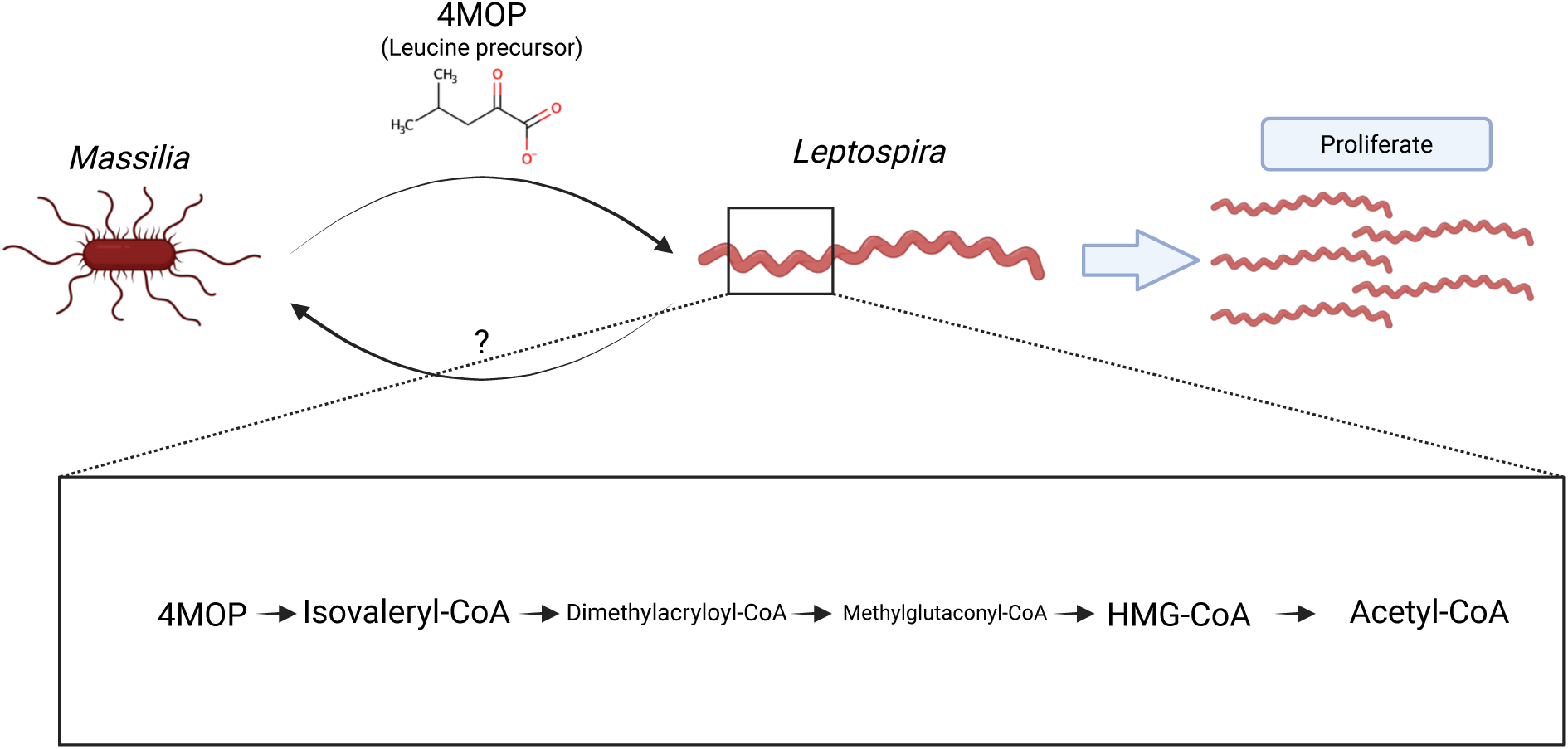
Proposed schematic model. This study proposes that *Leptospira* proliferates effectively by absorbing multiple metabolites secreted by *Massilia*, especially 4-methyl-2-oxopentanoate (4MOP), and converting them into acetyl-CoA. Whether metabolites released by *Leptospira* reciprocally affect *Massilia* growth remains to be determined in future studies. Created in BioRender. Ozuru, R. (2026) https://BioRender.com/5e8bt03

*Leptospira* displays unusually stringent nutritional requirements that are likely to shape both its cultivability and its environmental ecology (12, 18). A distinctive feature of pathogenic leptospires is their limited ability to use sugars as carbon sources, relying instead largely on beta oxidation of long-chain fatty acids for carbon and energy, consistent with the composition of standard media such as EMJH which supply fatty acids via Tween 80. In addition, *Leptospira* is a slow-growing organism *in vitro*, with reported doubling times commonly on the order of approximately 8 to 20 hours under laboratory conditions, and culture growth often requires prolonged incubation (19). These physiological constraints indicate that, in nutrient-limited environments such as soil and surface water, the growth and survival of *Leptospira* may be facilitated by diffusible metabolites released by neighboring microorganisms. In line with this view, isolation of pathogenic *Leptospira* from environmental samples remains challenging, in part because competing microbes and non-pathogenic leptospires can outgrow pathogenic strains during cultivation (20). Notably, supplementation of EMJH with 4MOP increased growth yield not only in saprophytic strains but also in pathogenic *Leptospira* in our experiments, highlighting the practical potential of metabolite-guided medium optimization. A clear next step will be to incorporate 4MOP and, related BCAA keto-acid intermediates when it is appropriate, into selective isolation workflows and systematically test whether recovery of pathogenic *Leptospira* from environmental samples can be improved without increasing background growth of competing taxa.

The growth-enhancing metabolites highlighted in this study, including 4MOP together with 3MOP and 3MOB, are BCAA-derived alpha keto-acids (21). This observation prompts discussion about how *Leptospira* accesses and metabolizes these compounds. Exogenous alpha keto-acids are typically transported by distinct classes of uptake systems from amino acids (22–25), and *Leptospira* may possess transport capacity for BCAA-derived keto-acids that may exceed the uptake of the free amino acids under the tested conditions. Also, keto-acids may enter the central metabolism pathway with fewer upstream processing steps than amino acids (26, 27), potentially reduce energy expenditure, and enable direct routing of carbon toward acetyl-CoA generating pathways. In contrast, amino acids uptake in bacteria is likely tightly regulated, and external amino acids may be preferentially channeled into anabolic demands or buffered by homeostatic controls rather than increasing net biomass accumulation (28, 29). These hypothesis that keto-acids are more efficiently incorporated into *Leptospira* than amino acids can be evaluated in a stepwise manner: (i) *in silico* identification of candidate alpha keto-acid transporters and relevant catabolic genes through genome-wide searches and comparative annotation, (ii) experimental assessment of their expression or induction under keto-acid supplementation, and (iii) functional tests using targeted gene disruption or chemical inhibition coupled to growth yield and uptake assays. Together, such analyses would clarify why BCAA-derived keto-acids, rather than the amino acids themselves, provide a more effective growth-promoting input for *Leptospira*.

In this work, we used genome-scale metabolic network reconstruction (GENRE) not as a stand-alone proof of mechanism, but as a principled framework for hypothesis generation and candidate prioritizations that complements high-throughput metabolite profiling (30). Genome-scale metabolic models have been widely applied across organisms (31–34) and contexts (35–41) including diverse microbial systems, linking genotype to phenotype, and guiding experimental design (42). Here, we have extended this modelling approach to a practical challenge in leptospiral biology, namely the cultivation difficulty of *Leptospira*, thereby assess the contribution of environmental and diffusible factors to growth yield. Metabolomics is powerful for identifying large numbers of compounds, but this breadth can complicate causal inference by producing a long list of candidates. By integrating metabolomics with a *Leptospira* GENRE, we were able to prioritize metabolites that are predicted to directly support biomass formation. Moreover, we show that draft metabolic models generated by automated pipelines such as Reconstructor are useful, provided that the model quality and limitations are transparently reported and evaluated (15, 43, 44). Moreover, transcriptome-informed contextualization using RIPTiDe (17) added an orthogonal layer of evidence by constraining feasible flux states under Msup conditions, offering a systems-level explanation consistent with increased utilization of leucine catabolic routes. Together, these results highlight the methodological importance of combining multi-omics data with GENRE-based modelling to move from broad chemical differences in conditioned media toward prioritized and testable growth-supporting mechanisms in microorganisms.

*Massilia* spp. are widely distributed and are frequently associated with soils and plant associated habitats (45, 46), making it plausible that their secreted metabolites contribute to shaping local chemical microenvironments *in situ* (3). In this context, an important open question is whether *Leptospira* and *Massilia* co-occur and interact in environmental reservoirs at spatial and temporal scales relevant to pathogen maintenance. As this work originated from a serendipitous observation during routine *Leptospira* cultivation (Fig. S1), which suggests co-existing property of these two organisms, it may add depth in a view of symbiosis. Fleming’s original report of penicillin similarly arose from an unexpected culture observation (47), underscoring how careful phenotypic monitoring in complex microbial settings can reveal previously unrecognized bioactive factors that shape microbial growth *in vitro*, and further, enable to construct realistic microbial interactions that can be examined.

We acknowledge several limitations in this study: First, although our data support the presence of growth-promoting BCAA-derived keto-acids in conditioned supernatants and their activity *in vitro*, we have not yet directly established the biosynthetic origin and secretion routes of these metabolites in *Massilia*. Because the concentrations of these keto-acids in Msup remain unknown, it is unclear whether the concentrations used in our supplementation experiments fall within physiologically and environmentally relevant ranges. To define the relevant biosynthetic genes and enzymes, and to quantify production and release under determined growth conditions, will be the next important steps. Second, the *Leptospira* GENRE used here is a draft reconstruction generated by an automated pipeline and has not undergone extensive manual and literature-based curation. The model captured multiple experimentally supported behaviors and proved it useful for metabolite prioritizations, illustrating the practical value of draft reconstructions when their scope and quality are transparently reported. Third, pathway-level interrogation highlighted a remaining knowledge gap on the *Leptospira* side: in our current annotation and modelling, a canonical enzyme for converting 4MOP-derived intermediates toward isovaleryl-CoA (48) could not be unambiguously identified (FIG 5).

Moreover, the transport mechanisms underlying keto-acid uptake remain unclear, and no dedicated transporter has been confidently identified in our current annotation. Resolving these gaps will require targeted annotation efforts and experimental validation, and it also raises the possibility that leptospires employ alternative enzymes or bypass routes (49) to channel keto-acid intermediates into acetyl-CoA generating metabolism.

## Materials and Methods

### Strains, media and reagents

The following strains were used in this study. *Leptospira*: *L. interrogans* serovar Manilae strain L495 (P1 clade); *L. johnsonii* strain E08 (P2 clade); *L. wholffii* strain YH112 (P2 clade); *L. kobayashii* strain E30 (S clade). *Massilia*: *Massilia* sp. strains NBRC 108631 (the National Institute of Technology and Evaluation,Tokyo, Japan). *Leptospira* strains were cultured in Ellighausen-McCullough-Johnson-Harris liquid medium (EMJH) consisting of Medium Base and Enrichment (279410 and 279510, respecrively; BD, Franklin Lakes, NJ), and *Massilia* was cultured in R2A liquid medium (Shiotani M.S. Co., Ltd., Hyogo, Japan). All cultures were incubated at 30°C. The following reagents were used: 4-methyl-2-oxopentanoate (4MOP; Sigma Aldrich, St Louis, MO), 3-methyl-2-oxopentanoate (3MOP; Sigma), 3-methyl-2-oxobutanoate (3MOB; Sigma), L-leucine (MP Biomedicals Germany GmbH, Eschwege, Germany), L-isoleucine (FUJIFILM Wako, Osaka, Japan), and L-valine (Nacalai Tesque, Inc., Kyoto, Japan).

### Preparation of *Massilia*-conditioned supernatant (Msup)

*Massilia* strains were cultured in R2A medium for 48 h. Cultures were centrifuged at 8000 × rpm for 5 min using a MX-205 (Tomy Seiko Co., Ltd., Tokyo, Japan) to isolate supernatants from pelleted cells. The resulting supernatants were passed through a 0.22 µm pore-size membrane filter Minisart NML (Sartorius AG, Göttingen, Germany) to generate *Massilia* culture supernatant (Msup). Msup was aliquoted and stored at −80°C until use. For each experiment, an aliquot was thawed on ice and used immediately; repeated freeze–thaw cycles were avoided.

### Gas Chromatography-Mass Spectrometry (GC-MS)/MS of Msup and downstream data analysis

Isolated culture supernatant (1 mL/sample) was freeze-dried for overnight at 45°C under 1 Torr with SpeedVac SPD140 DDA Vacuum concentrator that is connected to RVT5101 Refrigerated Vapor Trap and OFP400 Vacuum pump (Thermo Fisher Scientific KK, Tokyo, Japan). Dried materials were dissolved in 1 mL of extraction solution (methanol: water: chloroform = 2.5: 1: 1) with vigorous vortex. Samples were incubated at 37°C with 1,200 rpm agitation for 30 minutes in TAITEC MicroIncubator M-36 (TAITEC Corporation, Saitama, Japan). After centrifugation with 16,000 x g for 3 min at 4°C, 600 μL supernatant was mixed with 300 μL ultrapure water (214-01301, Fujifilm-Wako, Osaka, Japan). Samples were centrifuged with 16,000 x g for 3 min at 4°C, and 200 μL upper phase was freeze-dried. To the dried material, 100 μL of 20 mg/mL O-methylhydroxylamine hydrochloride in pyridine was added and incubated in MicroIncubator at 30°C, 1,200 rpm for 90 min (O-methylhydroxylamine hydrochloride M0343, Tokyo Chemical Industry Co., Ltd. (TCI), Tokyo, Japan, and pyridine 164-05312, Fujifilm-Wako). Then, 50 μL of N-methyl-N-trimethylsilyltrifluoroacetamide (MSTFA, M0672, TCI) was added and incubated further at 37°C, 1,200 rpm for 30 min. After brief centrifugation with 16,000 x g for 3 min at R.T., the supernatant was transferred to autosampler vial and used for GC-MS/MS (GCMS-TQ8050 NX, Shimadzu Corporation, Kyoto, Japan). The equipment was manipulated with GCMSsolution software Ver. 4.5 (Shimadzu) with n-alkane (C7 to C33) as qualitative retention time index standard (31080, Restek Japan, Tokyo, Japan), and samples were analyzed with multiple response monitoring (MRM) mode utilizing SmartDatabase (Shimadzu).

For downstream analyses, peak areas (Area) obtained from GC-MS/MS were used as quantitative measures. Where applicable, peak areas were normalized to the sample amount prior to multivariate analyses. For each set of samples, principal coordinates analysis (PCoA) was conducted in R (version 4.4.0) using the vegan package (version 2.6-8, (50)) based on a Bray–Curtis distance matrix. Group-level differences in overall metabolite profiles were evaluated by PERMANOVA using *adonis2* with 999 permutations. For univariate analyses, differences in individual metabolite abundances between groups were assessed using a two-sided Welch’s *t* test. Resulting *p* values were adjusted for multiple testing using the Benjamini–Hochberg method to control the false discovery rate (FDR). Metabolites meeting the significance threshold (adjusted *p* < 0.05) were visualized as heatmaps using the pheatmap package (version 1.0.12) with row-wise z-score applied.

### *Leptospira* proliferation confirmation experiment

Log-phase *Leptospira* cultures were enumerated using a Thoma cell counting chamber (Sunlead Glass Corp., Saitama, Japan) and diluted in EMJH medium to a final concentration of 1 × 10⁷ cells/mL. For Msup supplementation experiments, EMJH medium and R2A medium-diluted Msup were mixed at equal volumes for cultivation. Msup was diluted with fresh R2A medium to obtain final concentrations of 0%, 10%, 20%, 30%, 40%, and 50% in the assay. For metabolites supplementation experiments, reagents were added to EMJH medium diluted to 50% (vol/vol) with sterile water at the final concentrations indicated in the corresponding figure legends or in the Results section. Cultures were incubated at 30°C without shaking. Optical density at 450 nm (OD_450_) was measured every 24 h using a microplate reader (iMark; Bio-Rad Laboratories, Inc., Hercules, CA, USA). Growth curves were fitted from the OD_450_ time-series data using the R package growthcurver (version 0.3.1, (51)).

### Genome-scale metabolic network reconstruction (GENRE) analysis

GENRE of L495 was generated using Reconstructor (version 1.1.0, (15)). Protein sequence datasets were obtained from BV-BRC (https://www.bv-brc.org/) as amino acid annotation files for the genomes (Genome ID: 214675.23). For model reconstruction, medium conditions were set according to Table S1, and all other parameters were left at default settings. To evaluate the growth impact of candidate metabolites, we added exchange reactions for the uptake of each target compound to the reconstructed GENRE and quantified changes in predicted biomass production. Flux balance analysis was performed with the biomass reaction as the objective function. For selected compounds, uptake availability was systematically varied by adjusting the lower bound of the corresponding exchange reaction (negative flux indicating uptake) across a predefined range, and the resulting objective values (biomass flux) were recorded. All simulations were conducted in COBRApy (version 0.22.1, (52)) using the GLPK solver.

*In silico* supplementation was performed in COBRApy using the L495 genome-scale metabolic model. Candidate Msup-associated metabolites were tested individually by varying the carbon-equivalent exchange lower bound for each compound. Growth effects were quantified by flux balance analysis as dBiomass relative to the base medium control, and exchange and uptake fluxes were recorded to confirm utilization.

### Transcriptomics

*L. interrogans* L495 cells cultured for 48 hours under each condition (Fig. 4A) were harvested by centrifugation and total RNA was extracted from pellet using an RNA purification kit (Zymo Research Corp., Irvine, CA, USA). RNA-seq libraries were prepared using NEBNext rRNA Depletion (Bacteria) (New England Biolabs, Ipswich, MA), TruSeq Stranded Total RNA Library Prep Gold Kit (Illumina, Inc., San Diego, CA) and sequenced on an NovaSeq X (Illumina, Inc., San Diego, CA) to generate 151-bp paired-end reads. Protocol for rRNA removal using the NEBNext rRNA Depletion Kit (Bacteria), was from the “TruSeq Stranded Total RNA Reference Guide” (1000000040499 v00). Raw reads were subjected to quality control using FastQC (version 0.12.1) and adapter/quality trimming using BBDuk (version 39.08). Processed reads were aligned to the reference genome using STAR (version 2.7.11b), and gene expression was quantified using RSEM (version 1.3.3), using the *Leptospira interrogans* serovar Manilae strain L495 genome annotation from BV-BRC (Genome ID 214675.23) as the reference.

### Integration of transcriptome data and flux balance analysis

The L495 metabolic model and transcriptome data were integrated using RIPTiDe (version 3.4.81, (17)). A normalized RNA-seq count matrix (TPM) was used as input, therefore no additional normalization was applied within RIPTiDe. For each condition, RIPTiDe was used to generate a context-specific model, and flux sampling was performed with 1,000 samples per model using the RIPTiDe workflow. All computations were performed in Python using COBRApy and RIPTiDe with the solver set to GLPK.

### Statistical analysis

Statistical analyses were performed in R (version 4.4.0) using the vegan package (version 2.6-8) where applicable. Unless otherwise stated, statistical significance was defined as *p* < 0.05. For comparisons between two groups, the Wilcoxon rank-sum test was used unless otherwise specified in the relevant figure legends.

## Data Availability

All code used in this study, the generated genome-scale metabolic models, and the raw and processed data supporting the findings are available in a GitHub repository (https://github.com/O2U/Lepto_Massilia). RNA-seq data have been deposited in the NCBI Sequence Read Archive (SRA) under BioProject accession PRJDB40658 (DRR916665-DRR916673).

## Acknowledgement

We thank the members of the Department of Microbiology and Immunology, Faculty of Medicine, Fukuoka University for helpful discussions and technical support. We also thank Ms. Mizuki Yokono of Tottori Univ. Organization for Research Initiative and Promotion (ORIP) for performing GC-MS/MS. This work was supported by JSPS KAKENHI (Grant No. 24K11646 to M.Y. and 24K10225 to R.O.), AMED/NIH/CRDF (Grant No. JP23jk0210044 to F.O. and G.L.K.), AMED/NIH/ISTC (Grant No. JP25jk0210054 to R.O. and J.A.P.), and Takeda Science Foundation to R.O. The funders had no role in study design, data collection and analysis, decision to publish, or preparation of the manuscript.

## Supplemental materials

**Fig. S1 *Leptospira* growth promotion by *Massilia* culture supernatant (Msup).** Representative image from a preliminary experiment showing accelerated colony formation of *Leptospira* (arrow) in proximity to a culture supernatant from *Massilia* sp. Msup (round cup area) or less colonies at vicinity of the culture medium alone (arrowhead) on EMJH agar plate inoculated with *Leptospira* using Oxford cups. The original *Massilia* isolate used in this observation was not preserved, which motivated the present systematic study using multiple *Massilia* strains.

**Fig. S2 Comparison of *Leptospira* doubling times with or without Msup.** Doubling times were calculated from the growth curves shown in Fig. 1A. Within the present dataset, no statistically significant differences were observed in any comparison.

**Table S1 EMJH medium components for reconstruction of the *Leptospira* metabolic in this study.**

